# ConforFold Recovers Alternative Protein Conformations Beyond MSA Subsampling

**DOI:** 10.1101/2025.10.14.682366

**Authors:** Raulia Syrlybaeva, Eva-Maria Strauch

**Affiliations:** Department of Medicine, School of Medicine, Washington University in St. Louis, St. Louis, MO, United States; Department of Molecular Microbiology, Washington University in St. Louis, St. Louis, MO, United States; Department of Biochemistry and Biophysics, Washington University in St. Louis, St. Louis, MO, United States

## Abstract

Conformational changes underlie many aspects of protein function, yet current structure prediction tools remain limited in their ability to systematically sample structural ensembles. Here, we present **ConforPSSP** and **ConforFold**, a combined framework that integrates secondary-structure sampling into deep learning-based prediction to recover multiple protein conformational states. ConforPSSP employs a transformer model trained on multi-residue fragments to generate diverse 8-state protein secondary structure predictions (PSSPs), which are then used to condition a retrained OpenFold model (ConforFold). ConforFold achieved state-of-the-art performance in conformer recovery. On our test dataset of protein samples with two alternative conformations, it correctly identified both conformers in 84% of cases at TM-scores≥0.8, outperforming AlphaFlow (75.4%), which uses diffusion-based sampling, and Cfold, which relies on MSA clustering. Combining ConforFold with AlphaFlow further improved recovery rates while retaining the complementary strengths of both approaches. These results establish ConforFold as a broadly applicable framework for modeling structural ensembles. By explicitly integrating secondary structure it recovers conformations inaccessible to MSA-based subsampling or diffusion models, offering a new avenue for investigating conformational heterogeneity, mechanistic transitions, and the structural basis of protein function.

## Main

Conformational changes in proteins drive numerous biochemical activities. Identifying stable conformations is essential for understanding protein function and assembly mechanisms, providing molecular insights, controlling protein functions (or their suppression) ^1,2^, and developing novel therapeutics and subunit vaccines. Historically, molecular dynamics simulations have been used to model these fluctuations. Recently, advanced deep learning methods, notably AlphaFold2/3 ^3–5^ (AF2) and RoseTTAFold2^6^, have revolutionized protein structure prediction from amino acid sequences with high accuracy, independent of experimental data. However, these models, trained on static structures, are not optimized to reveal structural variations. A prevalent strategy to address this involves manipulating multiple sequence alignments (MSAs) and selecting templates to bias predictions toward different conformational states^7–10^. We reason that protein secondary structure prediction (PSSP) information could further guide models in predicting these diverse states.

Because protein conformations can change substantially with only a hinge motion, PSSPs need to be highly accurate and sensitive to detect structural ambiguities that guide the generation of structural ensembles. PSSPs can assist ensemble prediction either by enhancing template selection or by directly guiding protein folding predictions within the model. Our work focused on the latter approach: we retrained the open-source AF2 (OpenFold^11^) model, integrating PSSPs and using modified MSAs as input (Fig. 1).

**Figure 1.**
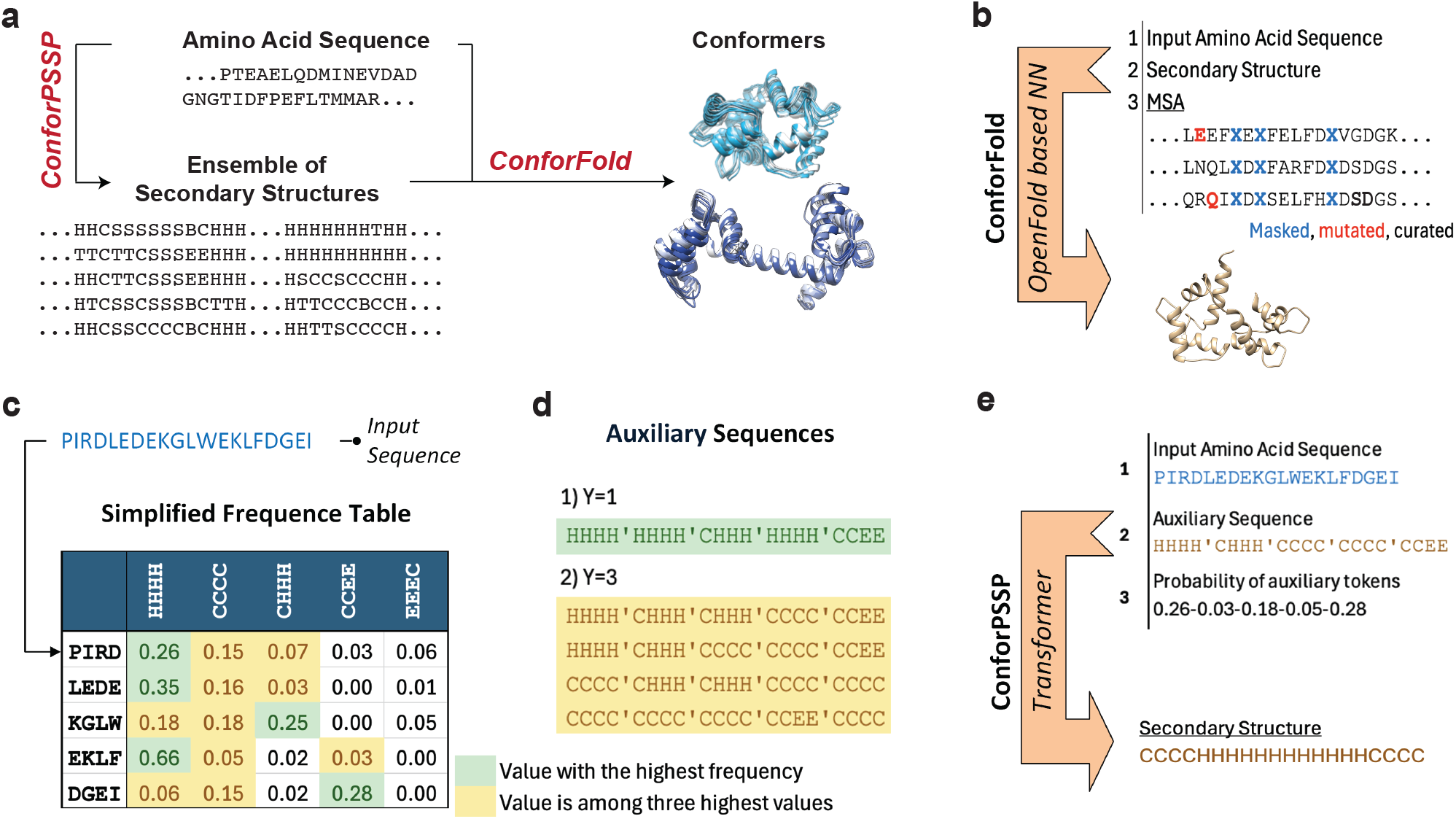
Details of the method. **(a)** Overview of a neural network framework for conformers generation based on predicting multiple PSSP for a query AAS using ConforPSSP at the first step, and folded conformations by means of ConforFold after. (**b)** Dataflow in ConforFold model. (**c)** Frequency table and its usage. (**d)** Auxiliary sequences based on the most probable (1) or randomly selected among the three topmost (2) PSSP chunks. (**e)** Dataflow in ConforPSSP model.

We developed ConforPSSP, a transformer model that predicts 8-state PSSP. According to the division method of the DSSP program^12,13^, the 8 states of protein secondary structure include G (helix), I (π-helix), H (α-helix), B (β-bridge), E (β-sheet), T (turn), S (bend), and C (coil). A core component of ConforPSSP is a frequency table built from amino acid sequences (AAS) in our training dataset and their corresponding protein secondary structures (PSS), which were segmented into 4-residue fragments. This normalized table quantifies the co-occurrence of unique 4-residue AAS and PSS fragments (Fig. 1c). For prediction, a query AAS is fragmented, and probable PSS fragments are randomly chosen for each AAS fragment based on the frequency table and assembled into an auxiliary sequence (Fig. 1d). The threshold number of the top PSS chunks used for selection is called *Y*. ConforPSSP’s output is then generated based on the input AAS, the auxiliary sequence, and PSS fragment probabilities retrieved from the table (Fig. 2e). Repeating this process produces diverse auxiliary sequences, resulting in different PSSP predictions. The innovation involves using statistical information from 4-residue fragments to produce multiple outputs, rather than relying solely on generative deep learning models.

**Figure 2.**
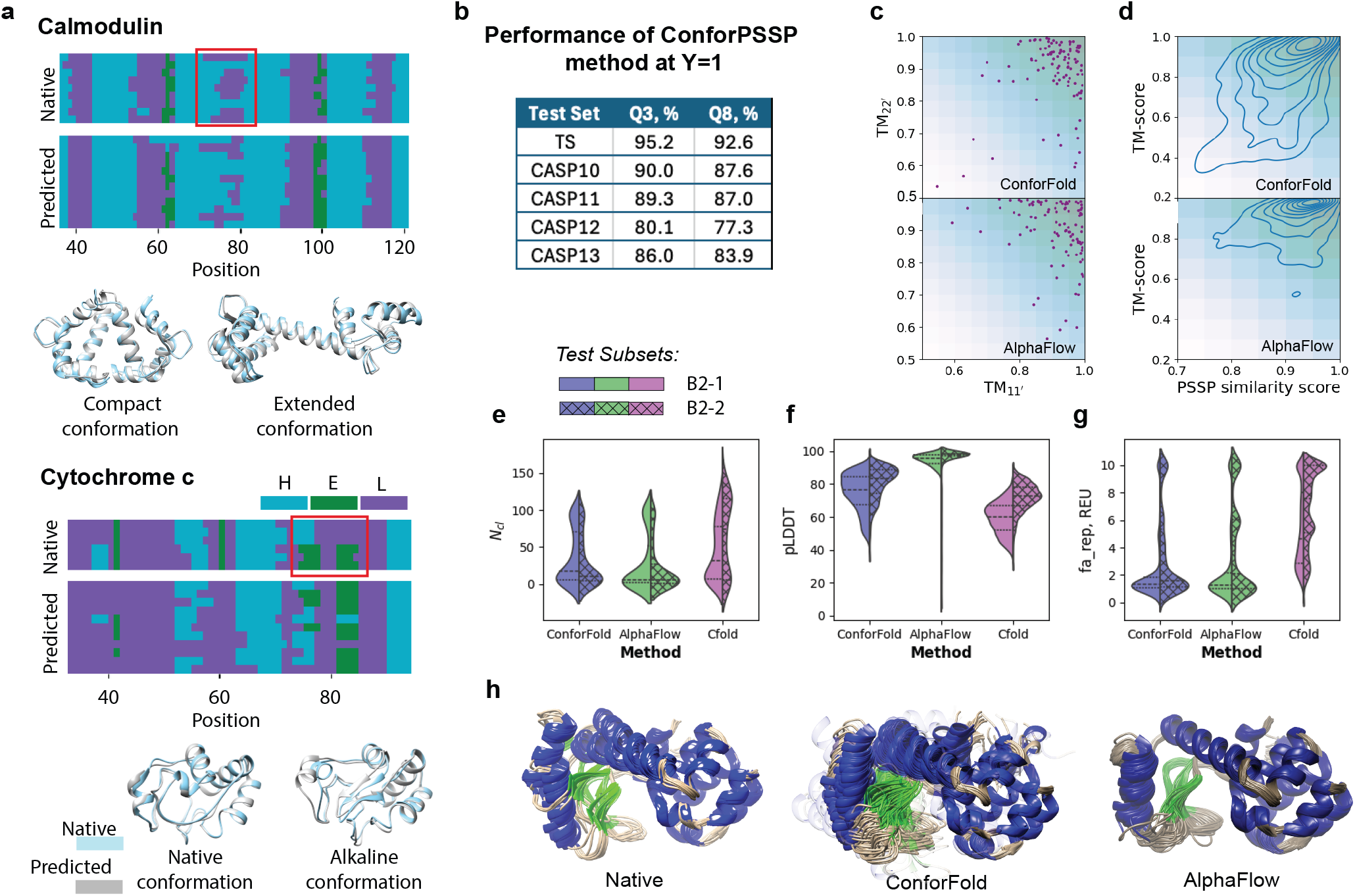
Performance of method. **a** Native and predicted PSSP and conformations of calmodulin and cytochrome. Input AAS for predictions are from 1CKK and 4Q5P models, respectively. Fragments of PSSP responsible for large conformational changes are in red frames. **b** Accuracy of ConforPSSP. **c** Comparison of performance of different methods on dataset B1, based on TM-scores for the most accurate predicted protein models corresponding to native (TM_11′_) and alternative (TM_22′_) conformations. **d** Comparison of performance of different methods based on distribution plot of correlation between PSSP similarity and TM-scores of all pairs of conformers across proteins of dataset B1. **eg** Performance of methods in terms of results diversity (**e**), the overall prediction confidence (**f**), and propensity for generating clashes (**g**) on datasets B2-1 and B2-2. **h** Comparison of native structures of lysozyme (PDB IDs 167l, 169l, 171l, 173l, 175l, 177l, 180l, 168l, 170l, 172l, 174l, 176l, 178l, 180l) with results of computational methods; results of ConforFold which differ from known conformers are depicted with transparency.

To evaluate our PSSP model, we used a test subset (TS) and CASP datasets^14^ (Fig. 2b). The accuracy was measured at Y=1 in terms of 3-state PSSP (Q3), 8-state PSSP (Q8), and segment overlap scores are reported as well (Fig. 2b, Supplementary table 1). Our prediction accuracy on TS dataset ranges from 92% to 95% across all metrics.

To sensitize one of the leading structure prediction algorithms, we retrained OpenFold. The resulting model was named ConforFold. Structure prediction algorithms typically rely on multiple sequence alignments (MSA)^7–10,15–17^, based on the idea that the evolutionary history of proteins reveals structurally and functionally important residues through their co-evolution. Popular methods for conformational prediction include using shallower MSAs^7,10,17^ or randomly masking MSA positions to increase uncertainty. We therefore reduced the number of MSAs used and masked 40% of the MSA positions randomly, inspired by the SPEACH_AF model^16^. Finally, we added “noise” to the MSA by randomly mutating 6% of positions.

Overall, these changes not only increased structural diversity but also improved the model’s responsiveness to PSSP conditioning—an essential adjustment since training started from OpenFold’s pre-trained weights. To encourage greater use of secondary structure input, we also masked the query AAS in the same positions as in MSA during training (but not during testing). Since neural networks (NN) perform best on samples typical for the training dataset, we used a unified input MSA: we replaced gaps with corresponding residues from a query sequence, and if the number of available MSA sequences was low, we increased their number by duplicating random MSAs for predictions.

We assessed ConforFold, used with PSSP ensembles from ConforPSSP, on a test set B1 of 62 proteins, containing samples with known two conformers. This set, largely overlapping with a prior one from Cfold method^10^, was augmented with examples of fold switches and some other. Predicted PSSPs were fed to ConforFold to predict their structures. Examples of the test cases are performed in Fig. 2a. Fold-switching proteins like Calmodulin (CaM) and Cytochrome C (Cyt c) are difficult to model (Fig. 2a). CaM, a well-studied example, shifts from a compact, Ca2+-free state (disordered central linker) to an extended, Ca2+-saturated state where the linker forms a rigid ‘central helix’ ^18^. The PSSP contain samples corresponding for both states (Fig. 2a), though the compact form was more common. Cytochrome C undergoes an ‘alkaline transition’ to alternative conformations, driven by a Met80 to Lys substitution that rearranges loop 70-85. This alters the most flexible loop, which is a β-hairpin in the native state at neutral pH. PSSPs from the alkaline Cyt c model (4Q5P_A) exhibited a mixture of native and alkaline types, with a slight preference for the alkaline form. However, PSSPs from the native Cyt c model’s sequence (3NWV_A) only yielded native-like secondary structures, likely due to the Met80 difference (Supplementary table 2).

We generated 100 conformers for each test sample of dataset B1 and computed TM-scores by comparing them against native structures. For each amino acid sequence (AAS), we selected the output conformers that most closely matched the folding of query AAS and its alternative conformation (denoted scores TM_11′_ and TM_22′_, respectively). As a benchmark, we compared performance against AlphaFlow, a diffusion-based model that delivers highly realistic structures and offers a balance of precision and diversity compared to AlphaFold with MSA subsampling ^19^.

ConforFold successfully generated both conformers in 84% of cases (104/124) at the 0.8 TM-score threshold. By comparison, AlphaFlow recovered both conformers in 75.4% of cases— slightly less accurate than ConforFold—although ConforFold produced more outliers (Fig. 2c). When AlphaFlow was supplied with ConforFold outputs as templates (AlphaFlow+ConforFold), both conformers were observed in 83% of cases (Supplementary Fig. 6–7). For reference, Cfold, a deep learning approach based on MSA clustering, identified just over 50% of nonredundant alternative conformations in their dataset^10^.

Finally, we tested the effect of random mutation “noise” on ConforFold’s performance. Removing the 6% mutation noise significantly reduced its accuracy, with successful identification of both conformations dropping to 56% (Supplementary Fig. 5).

The key strength of our method lies in its ability to identify previously uncharacterized protein conformations. To assess this, we evaluated ConforFold using datasets B2-1 and B2-2, which contain proteins with only one known conformer and poor-quality MSAs that typically challenge existing prediction methods. We benchmarked ConforFold against AlphaFlow and Cfold. For each sequence, 100 structural models were generated by all methods except Cfold, which produced between 140 and 150 models on average. Predicted structures were subsequently relaxed, clustered at a TM-score ≥ 0.8, and the highest pLDDT model from each cluster was used for evaluation.

ConforFold consistently outperformed Cfold in model quality, as reflected by higher pLDDT distributions (Fig. 2f) and lower repulsion energies during folding (Fig. 2g). Compared with AlphaFlow, ConforFold achieved greater conformational diversity (Fig. 2e) and produced models with the fewest steric clashes among the three methods (Fig. 2g). AlphaFlow, however, remained the top performer in terms of absolute pLDDT scores.

Across all three test sets, methods based on MSA subsampling, such as Cfold and ConforFold, demonstrated the ability to generate diverse structural ensembles. Importantly, ConforFold extended beyond the capabilities of simple MSA subsampling by incorporating secondary structure information, enabling it to recover conformers that remain inaccessible to standard approaches. This advantage translated into greater structural accuracy and precision, with ConforFold consistently outperforming Cfold in terms of both pLDDT distributions and folding energetics (Fig. 2f–g).

AlphaFlow, in contract, produced less diversity than methods based on MSA subsampling, as it was mentioned; 91.5% of conformers generated by AlphaFlow have pairwise TM-scores higher than 0.8 across dataset B1 according to our calculations (Fig. 2d), and, therefore, they can be considered as the same folding according to our threshold. Comparison of output structures depicted in Fig. 2h shows that AlphaFlow tends to generate outputs with very similar, homogenous, regions. While AlphaFlow was able to predict structures with different secondary structural sequences within its predictions, these modifications often did not correspond to substantial changes in overall folding. By comparison, ConforFold captured a more prominent relationship between secondary structure diversity (PSSP) and TM-score variation, reflecting its heightened sensitivity to conformational heterogeneity (Fig. 2d).

Taken together, these findings highlight complementary strengths of the methods: AlphaFlow excels at producing consistently high-confidence structures, whereas ConforFold provides a more nuanced exploration of conformational space, uncovering alternative states that would otherwise be missed by MSA-driven or diffusion-only strategies.

In this conclusion, we introduce a novel framework for sampling protein conformations that explicitly integrates secondary structure dynamics/variations into deep learning-based structure prediction. At its core, the method leverages a deep learning model capable of capturing shifts in secondary structure, which is further enhanced by a statistical PSSP strategy applied to multi-residue fragments. By combining this approach with a retrained state-of-the-art structure prediction algorithm, our system is able to generate multiple high-quality conformations for a given sequence. This capability establishes the framework as a broadly applicable platform for structural ensemble modeling, offering new opportunities to investigate conformational heterogeneity, mechanistic transitions, and the structural underpinnings of protein function.

Source code for the ConforPSSP and ConforFold models, trained weights and inference script are available under an open-source license at https://github.com/strauchlab/Confor-PSSP and https://github.com/strauchlab/ConforFold.

## Methods

### 1. Details of ConforPSSP

#### 1.1. The frequency table

We segmented all amino acid sequences (AAS) in our dataset (Supplementary Figure 10), along with their corresponding secondary structure annotations, into overlapping 4-residue fragments. These were used to construct a database, referred to as the frequency table, which records the normalized occurrence of each 4-mer AAS fragment associated with a specific PSS label (Figure 1c). Conceptually, this frequency table functions similarly to attention matrices in transformer neural networks, capturing the strength of the relationship between input tokens (AAS fragments) and output tokens (PSS annotations).

To capture more information, the frequency table incorporates not only a target AAS fragment but also its immediate neighboring fragments. The data structure is implemented as a Python dictionary, where each key represents a central AAS fragment, and the corresponding value is a nested dictionary. In these nested dictionaries, the keys represent adjacent AAS fragments (preceding and succeeding the central one), and the values are dictionaries mapping PSS tokens to their observed frequencies in that local sequence context.

To account for varying representation sizes of different AAS fragment clusters and to ensure fair contribution across the dataset, frequency values F are normalized using the following formula:

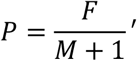

Here, *M* is defined as the base-2 logarithm of the product of the sample counts from all clusters associated with the target AAS fragment, as shown in Supplementary Figure 10b, prior to any down sampling. The value *P* represents the raw frequency used within the frequency table. The resulting database is saved in the file “*freq_table.json*” as part of the ConforPSSP framework.

#### 1.2. Inputs

Each query amino acid sequence (AAS) is decomposed into not overlapping 4-residue fragments, which are represented as AAS tokens. While the total number of possible unique 4-mer fragments composed of all 20 amino acids is quite large, we observed a slight improvement in PSSP accuracy when certain amino acids were substituted with mutagenically similar ones (with similar chemical properties). As a result, throughout the entire ConforPSSP pipeline, including construction of the frequency table, we apply the amino acid substitutions listed in Supplementary Table 3. After these substitutions, the total number of unique 4-mer AAS tokens is reduced to 103,849.

The maximum allowed length for an AAS is set to 800 residues. Sequences exceeding this length are truncated, while shorter sequences are padded with zeros. A special start token (“1”) is inserted at the beginning of each sequence, and an end token (“2”) is appended at the end. Any alignment gaps within the AAS are encoded using a dedicated gap token (“3”), and sequences are further padded with unknown residue tokens (“4”) as needed to ensure their total length is divisible by four. During training, each sequence is processed four times, with each pass using a different starting position among the first four residues to generate distinct samples (see Supplementary Table 4).

A second input to the model is a frequency-based sequence, constructed from the AAS tokens. For each 4-mer AAS fragment, we randomly select a corresponding PSS fragment from among the most probable PSS tokens, based on the frequency table, to form an auxiliary predicted secondary structure sequence. The total number of unique 8-state, 4-residue PSS tokens used for both input generation and model output is 2,370. Additionally, an auxiliary NN input consists of an array of probability values *P*, representing the likelihoods of the selected PSSP tokens derived from the frequency table.

#### 1.3. Outputs

Final predictions are generated using three inputs: AAS, the corresponding frequency-based sequence, and the associated probability array *P* (Figure 1c). By default, we evaluate four versions of each input AAS, each offset by one residue to account for different starting positions. Among these, the predicted PSSP sequence with the highest confidence score is selected as the final output. Confidence is assessed by summing the softmax values from the output layer across the sequence, treating these values as token-level probabilities.

To enhance diversity in the predicted secondary structures, this procedure is repeated multiple times per sample. Each iteration generates a new frequency-based sequence by sampling from the frequency table, thereby producing a different candidate PSSP output.

#### 1.4. Architecture and Training of ConforPSSP

The ConforPSSP model is based on a standard Transformer (“vanilla”) architecture. The encoder receives the amino acid sequence (AAS) as input, while the decoder takes the corresponding frequency-based sequence. Additionally, the probability array *P*, derived from the frequency table, is passed through a fully connected layer to match the dimensionality of the auxiliary sequence embeddings. The resulting vector is then summed with the auxiliary sequence embeddings prior to the addition of positional encodings in the decoder.

Training was performed with a sequence length parameter Y=7, across 10 epochs, repeated 15 times. From these, two models were selected based on manual evaluation, prioritizing both the correctness and diversity of predicted PSSP outputs.

The training loss is a categorical cross-entropy function composed of three components: The cross-entropy margin between predicted and true PSSP sequences.

Prediction accuracy for tokens where the auxiliary sequence does not match the ground truth, enforcing correction of uncertain regions.

The cross-entropy margin between the auxiliary sequence and the ground truth PSSP, encouraging alignment and promoting diversity.

To further guide the model, 20% of the ground truth PSSP tokens were intentionally included in the auxiliary sequences. Additionally, to prevent overfitting in cases where an AAS token has only one corresponding PSSP label in the frequency table, a random PSSP token was substituted in 20% of such instances during training.

The model was implemented using the TensorFlow framework.

### 2. Details of ConforFold

ConforFold is a retrained OpenFold’s model 3, which does not use templates for predictions, starting with its pre-trained weights. ConforFold does not utilize structural templates for prediction, a decision made to accommodate computational constraints. A key distinguishing feature of ConforFold is its incorporation of predicted secondary structure information as an additional input.

#### 2.1. Inputs of ConforFold

We initialized our model using pre-trained weights from OpenFold that already corresponded to a local minimum of the loss function. As a result, direct integration of secondary structure signals required deliberate intervention to ensure their influence on predictions. To achieve this, we intentionally degraded the reliability of the original OpenFold inputs, AAS and MSAs, so that the model would rely more heavily on the provided secondary structure information. This strategy aligns with established approaches for generating conformational diversity in folding networks via MSA perturbation, as discussed in the main text.

Specifically, we randomly masked 40% of positions in both the AAS and MSA inputs during training. During inference, only the MSA remains masked, while the AAS is provided in full. Additionally, we reduced the depth of MSAs by randomly sampling a subset of homologous sequences, which increases uncertainty and encourages conformational sampling. To further introduce variability, 6% of MSA positions were randomly mutated. The PSSP inputs, by contrast, were used without any manipulation.

To enhance model performance on the test set, we curated the MSAs provided during inference. Neural networks tend to perform best on inputs that resemble their training data; thus, we standardized MSA profiles to more closely align with those seen during training. For samples with extremely shallow MSAs, we duplicated existing sequences to increase MSA depth and replaced gaps with the corresponding residues from the query sequence. This curation procedure significantly reduced structural clashes in predicted models, particularly in the B2-1 and B2-2 datasets.

#### 2.2. Difference between architectures of ConforFold and OpenFold

8-State PSSP was provided to ConforFold as an additional input. Input AAS and PSSP are converted to one-hot vectors and concatenated together. The result is processed in the main “Input Embedder” block, instead of just AAS one-hot vectors.

#### 2.3. Training of ConforFold

The training was conducted in two phases through 5 epochs each. During the initial training phase, sequences were cropped to 256 residues, MSA depth was capped at 128 and extra MSA depth was capped at 1024. During fine-tuning, sequences were cropped to 384 residues; values corresponding to MSA parameters were not changed. The model was retrained using “finetuning_no_templ” model configuration both times with changes mentioned above.

### 3. Main Dataset

The dataset used to train ConforPSSP and ConforFold consists of around 140,000 entries from RCSB PDB as of August 2024 (Supplementary Figure 10). Protein sequences of RCSB PDB were selected according to criteria presented in Supplementary Figure 10a. The structures were complemented by their secondary structure information, PSSP were obtained using DSSP software and its database ^12,13^. Pre-computed MSA to train ConforFold were obtained from the OpenFold dataset, which is available at the Registry of Open Data on AWS (RODA)^20,21^. The dataset was organized in clusters (Supplementary Figure 10b) based on sequence cluster IDs provided in RCSB PDB and similarity of secondary structures calculated using Biopython.

#### 3.1. ConforPSSP dataset

Clusters of the main dataset at 70% sequence identity were split into training, validation and test subsets (Supplementary Figure 10c). We downsized big clusters by random representative selection during each epoch of training (Supplementary Figure 10d) to avoid bias toward populated clusters. Therefore, composition of the training data is different for each epoch of training process.

Beside the test set (TS) of the method, CASP datasets (Supplementary Table 5), namely CASP2010, CASP2011, CASP2012, CASP2013^14^, were used for benchmarking as well. Additional test set B3-1 with AAS sequence clusters at 50% identity consisting of 5 or more entries was used to check performance of the methods on a broad scope of structures; each cluster was downsized to 5 samples by random selection, total number of clusters is 14819.

#### 3.2. ConforFold dataset

Samples that were the sole representatives in AAS clusters at 70% sequence identity were excluded from the primary dataset. These excluded instances were used to construct a validation set of 2,000 sequences and the test sets B2-1 and B2-2 (see Supplementary Table 7). Cluster downsizing during training followed the same procedure as described for ConforPSSP.

The B2-1 and B2-2 test sets were designed to evaluate the model’s ability to predict unknown alternative conformations. B2-1 includes proteins for which OpenFold provides shallow MSAs, while B2-2 includes proteins with deeper but highly gapped MSA alignments. Each of these sets contains 200 protein samples.

The B1 dataset (Supplementary Table 6) was constructed primarily from the Cfold dataset ^10^ to assess the model’s capability in multi-state design, specifically for proteins with known alternative conformations. Each B1 entry includes two structural conformations, ordered alphabetically. Most B1 proteins were retained in the training set because the B1 dataset was significantly modified after training was completed. Initially, it included only entries with pairwise PSSP similarity below 85%, but later all entries from Cfold were incorporated.

We believe this overlap does not significantly bias the results due to several factors:

- In most cases, the query sequences corresponding to alternative conformations differ, making it inherently difficult to recover both states from the same AAS input;
- Only one representative was sampled per cluster at 90% sequence identity per epoch, and training was limited to 10 epochs. This meant that most samples in dense clusters were not utilized during training.
- The training dataset itself was noisy (masked and mutated), further reducing the likelihood of overfitting.

The B3-2 dataset was used to assess model performance on a diverse set of protein structures from the RCSB PDB. The main dataset (Supplementary Figure 10a) was clustered at 50% sequence identity using RCSB-provided cluster IDs. Clusters containing more than 50 sequences were selected, and 5 samples were randomly chosen from each for inclusion in B3-2, resulting in a total of 1,015 clusters.

### 4. Testing of the methods

#### 4.1. Usage of ConforPSSP on test subsets

A single secondary structure was predicted for each sample of TS, CASP datasets, and B3-1 dataset. Predictions for B3-1 were conducted at Y=15; in other cases, the results were obtained at Y=1 and Y=15. The samples with the highest and lowest Q8 prediction accuracy were identified for each cluster in B3-1 dataset.

#### 4.2. Multi-state predictions for dataset B1 using ConforFold

Secondary structure sequences were generated using two ConforPSSP models, each model produced 50 sequences at Y=15 (100 in total). Each secondary structure was used for prediction of a single output using ConforFold. The outputs were not relaxed.

#### 4.3. Multi-state predictions for dataset B1 using AlphaFlow

Predictions were performed using the base version of AlphaFlow-PDB model. 100 Outputs were generated for each input sequence. We could not do calculations for all inputs from dataset B1 using our gpu (NVIDIA L40S), therefore, the total number of inputs is 106 instead of 124. Rate of samples for which both conformations were recovered is calculated here and in section 4.4 against the reduced number of inputs.

#### 4.4. Multi-state predictions for dataset B1 using AlphaFlow+ConforFold

Predictions were performed using the base version of AlphaFlow-MD+Templates model. Each output of ConforFold (section 4.2) was provided to AlphaFlow as a single template once, resulting in 100 predictions. We were able to make calculations for 106 sequences.

#### 4.5. Multi-state predictions for datasets B2-1 and B2-2 using ConforFold

Secondary structure sequences were generated using two ConforPSSP models, each model produced 50 sequences at Y=15 (100 in total). Each secondary structure was used for the prediction of a single output conformation using ConforFold. The outputs were relaxed by means of Amber relaxation module of OpenFold. Conformations corresponding to the native structure (TM-score ≥0.8) were excluded to identify unknown conformers only. The remaining structures were clustered based on TM-score with a threshold set to 0.8, and a sample the highest pLDDT was selected from each cluster.

#### 4.6. Multi-state predictions for datasets B2-1 and B2-2 using Cfold

Conformational states of proteins were generated using 20 samples and five recycles per cluster size; the total number of generated outputs per protein is around 140. The structures, for which the repulsive Rosetta score term (fa_rep) was higher than 7,000 (scorefunction ref2015) for any residue, were discarded. Remaining outputs were relaxed by means of Amber relaxation module of OpenFold. Conformations corresponding to the native structure (TM-score ≥0.8) were excluded after. The outputs were clustered based on TM-score with a threshold set to 0.8, and the highest pLDDT was selected from each cluster.

#### 4.6. Multi-state predictions for datasets B2-1 and B2-2 using AlphaFlow

Predictions were performed according to the procedure described in section 4.3. Predicted outputs were processed as described in 4.5.

#### 4.7. Multi-state predictions for datasets B2-1 and B2-2 using AlphaFlow+ConforFold

Predictions were performed according to the procedure described in section 4.4 based on outputs of ConforFold (section 4.5). Predicted outputs were processed as described in 4.5.

#### 4.8. Folding of samples from dataset B3-2 using ConforFold

A single folded structure was predicted for each sample of the dataset using the native secondary structure. The samples with the highest and lowest TM-score between native and predicted structures were identified for each cluster.

## Supplementary information

### Supplementary Figures

**Supplementary Figure 1.**
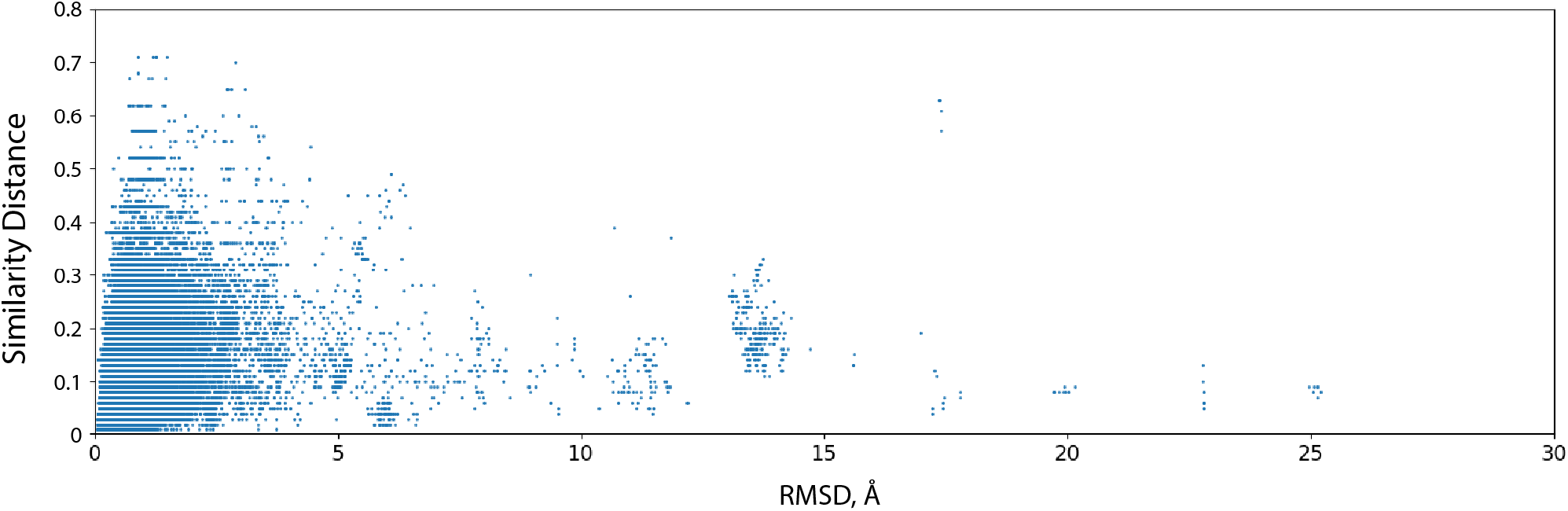
Secondary structure similarity between pairs of protein conformations obtained from RCSB Protein Data Bank and corresponding RMSD between their Ca atoms

**Supplementary Figure 2.**
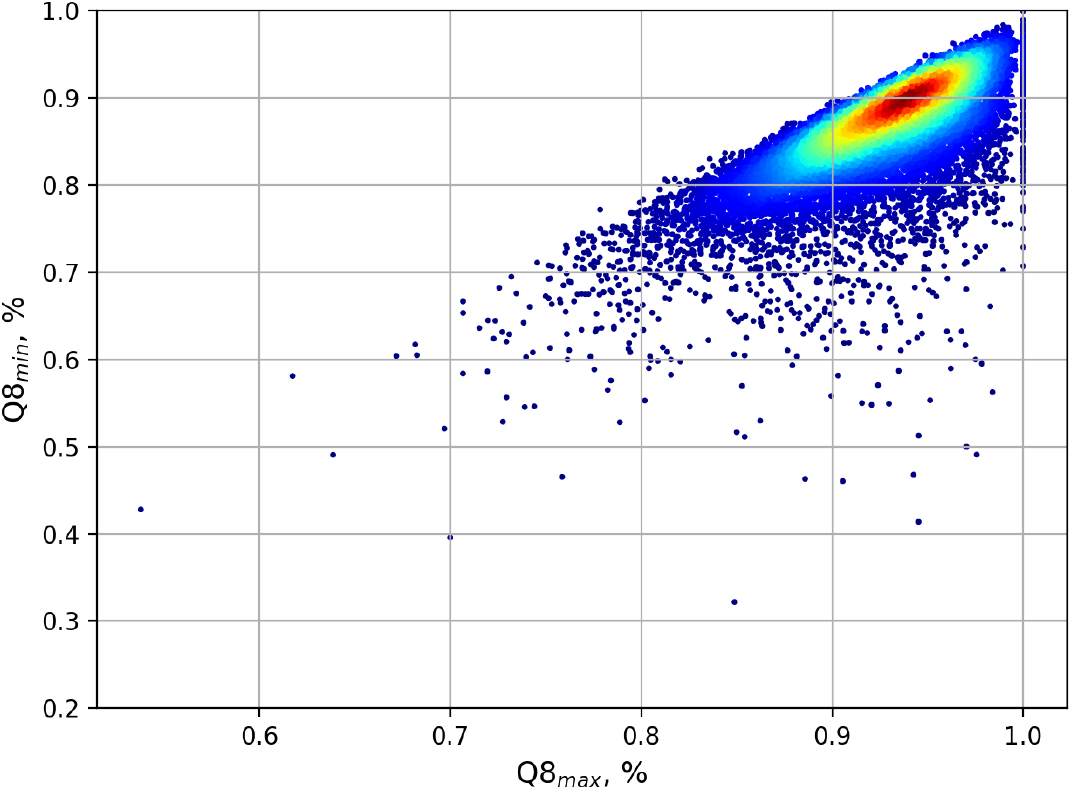
The highest (Q8_max_) and lowest (Q8_min_) accuracy values in terms of Q8 obtained after PSSP generation for dataset B3-1 (SI-2.1.5, SI-2.4.1).

**Supplementary Figure 3.**
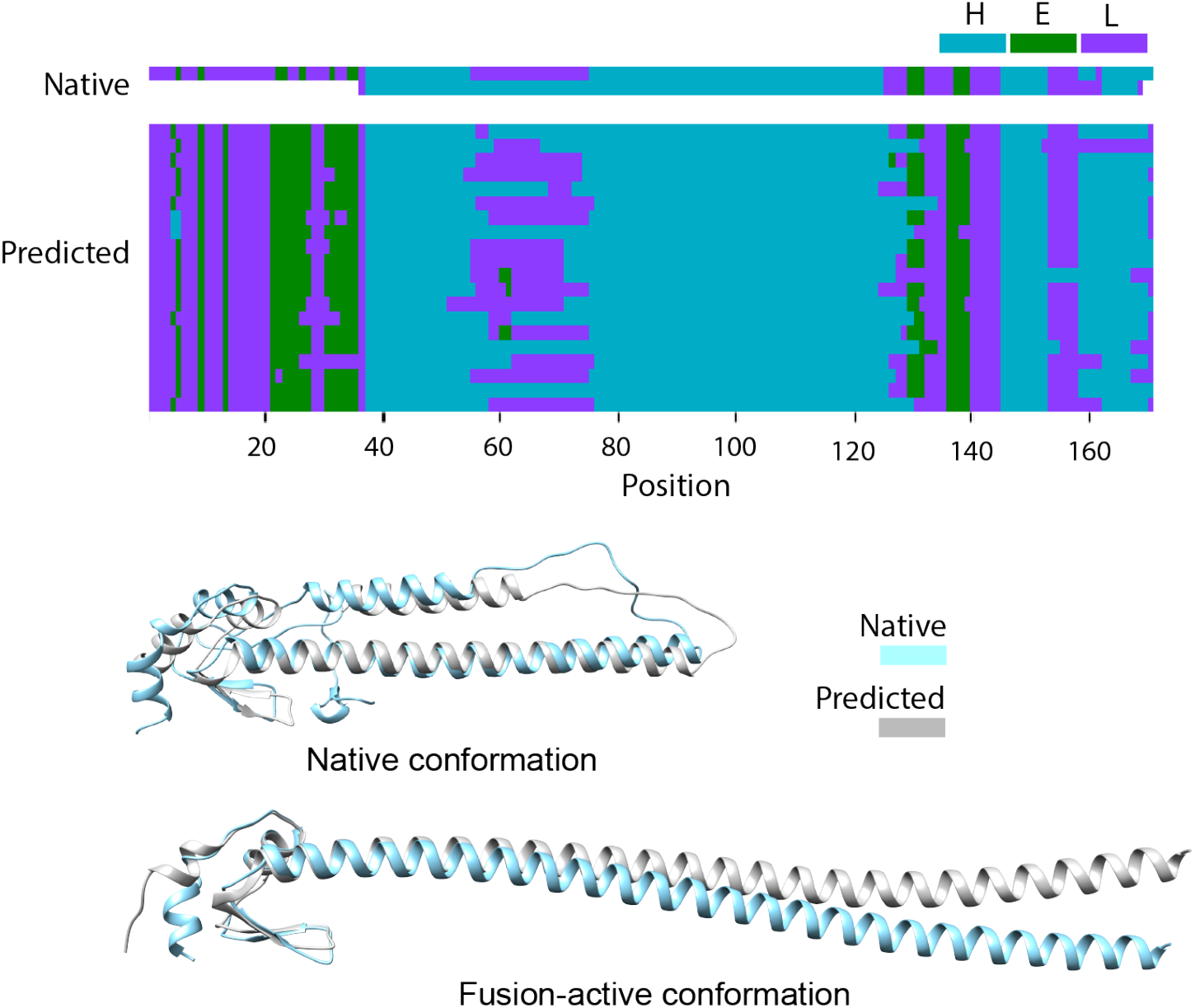
Native and predicted conformations of influenza hemagglutinin. Predictions are based on AAS from 6Y5H_B.

**Supplementary Figure 4.**
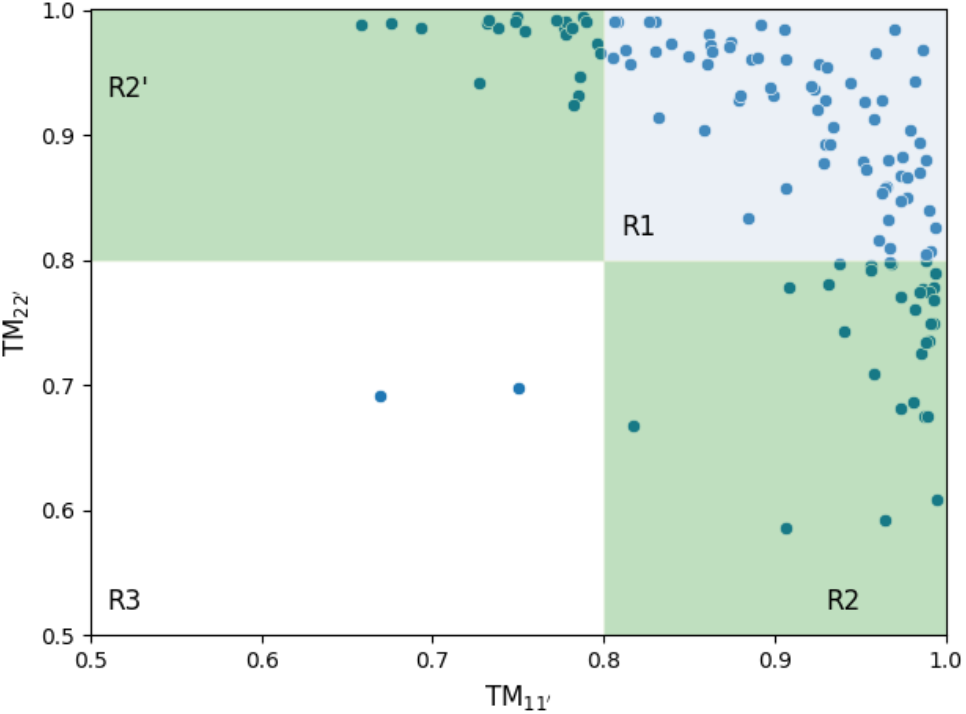
Distribution of TM-scores for the most accurate predicted models corresponding to native (TM_11′_) and alternative (TM_22′_) conformations, generated for dataset B1 by ConforFold without MSA noising.

**Supplementary Figure 5.**
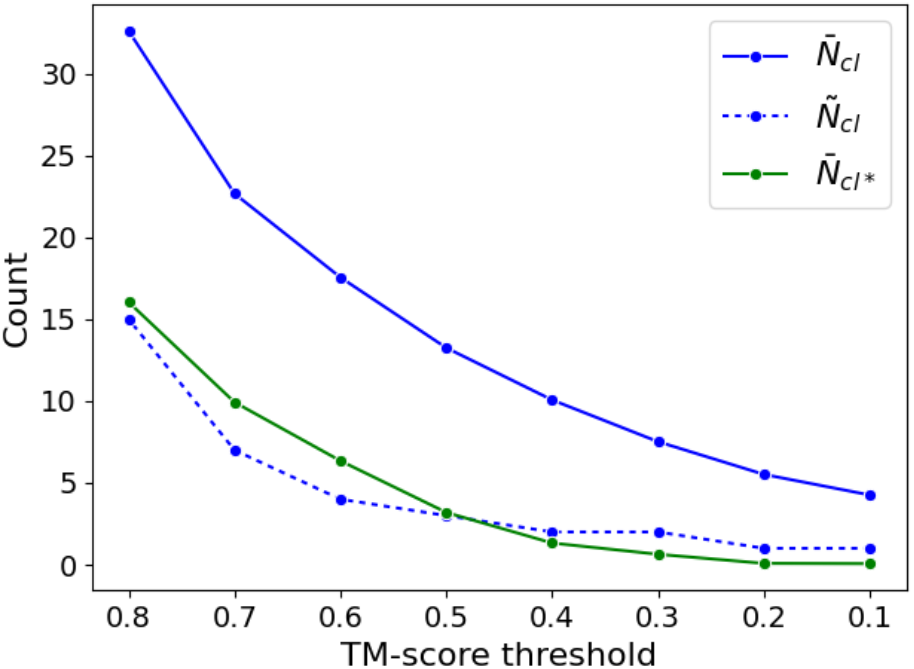
Number of structural clusters per input obtained over datasets B2-1 and B2-2 at different TM-score similarity thresholds: average and median number (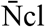and Ñ cl, respectively), average number of clusters with TM-scores < 0.8 relative known conformers 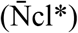.

**Supplementary Figure 6.**
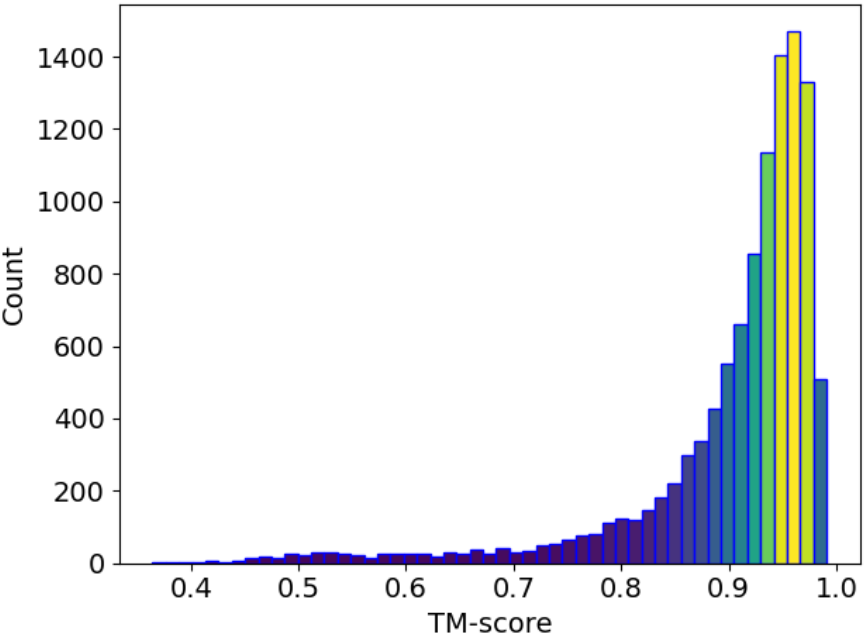
TM-score between templates and outputs of AlphaFlow+ConforFold approach.

**Supplementary Figure 7.**
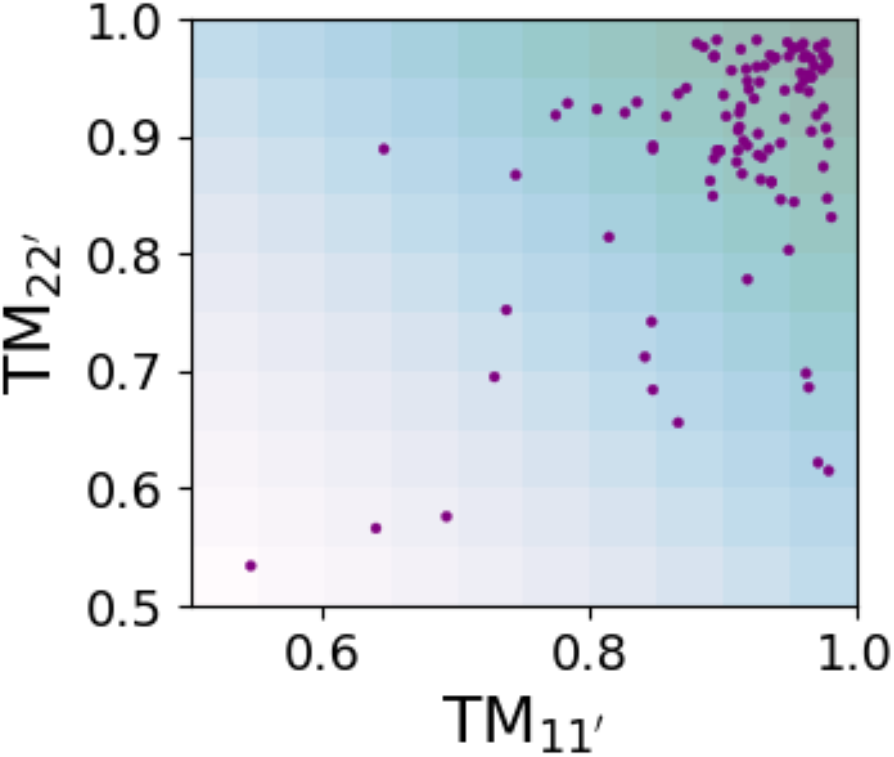
Performance of AlphaFlow+ConforFold approach on dataset B1 based on TM-scores for the most accurate predicted protein models corresponding to native (TM_11′_) and alternative (TM_22′_) conformations.

**Supplementary Figure 8.**
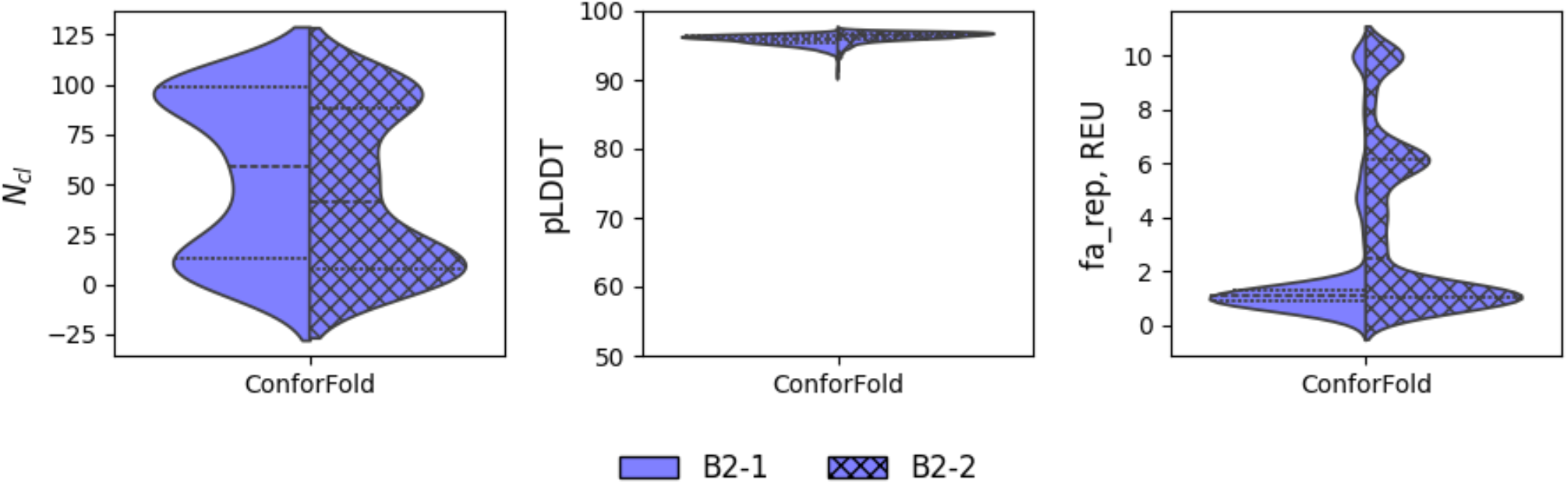
Performance of AlphaFlow+ConforFold approach in terms of results diversity (N_cl_), the overall prediction confidence (pLDDT), and propensity for generating clashes (fa_rep) on datasets B2-1 and B2-2.

**Supplementary Figure 9.**
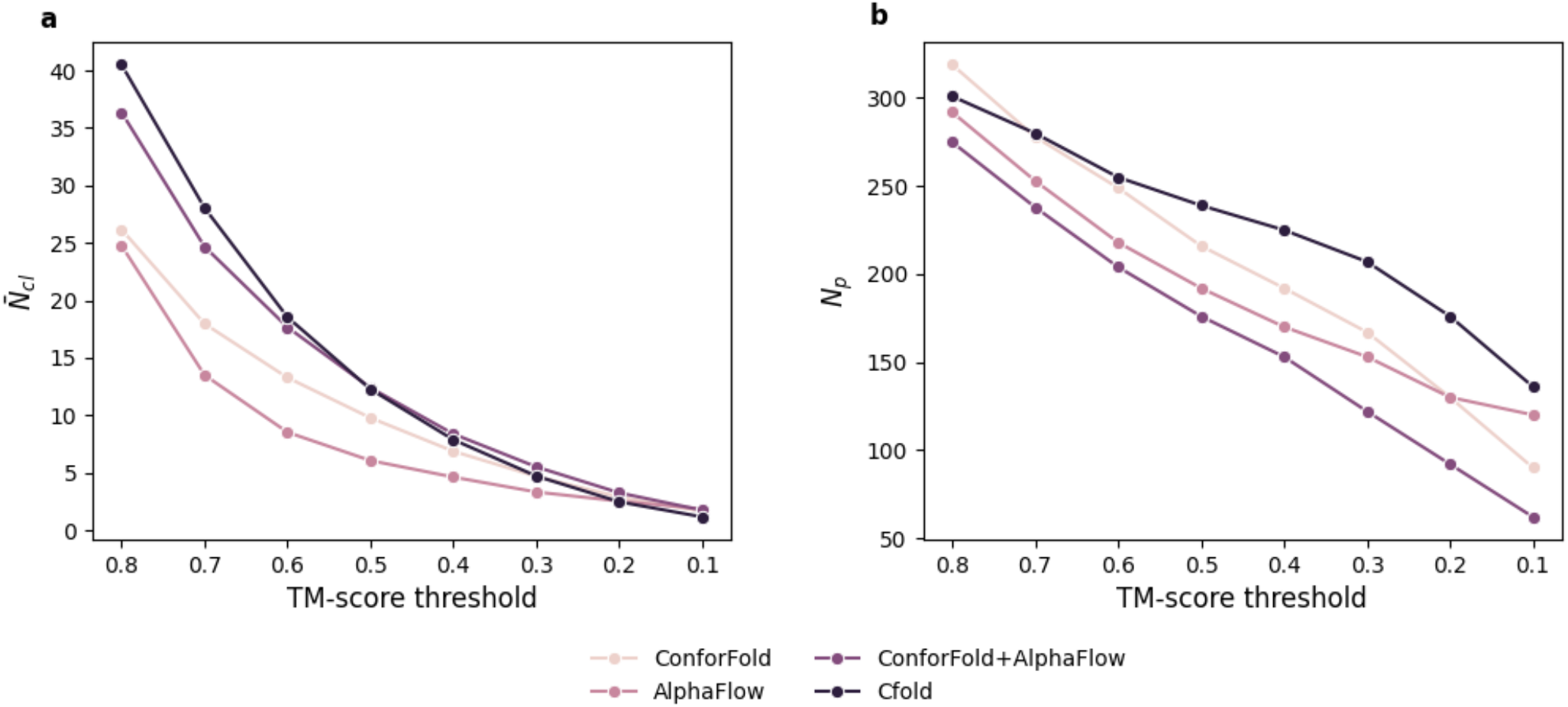
Performance of different methods on datasets B2-1 and B2-2 regarding diversity of outputs (**a**) and number of proteins for which more than one conformer is predicted (**b**) depending on applied TM-score threshold.

**Supplementary Figure 10.**
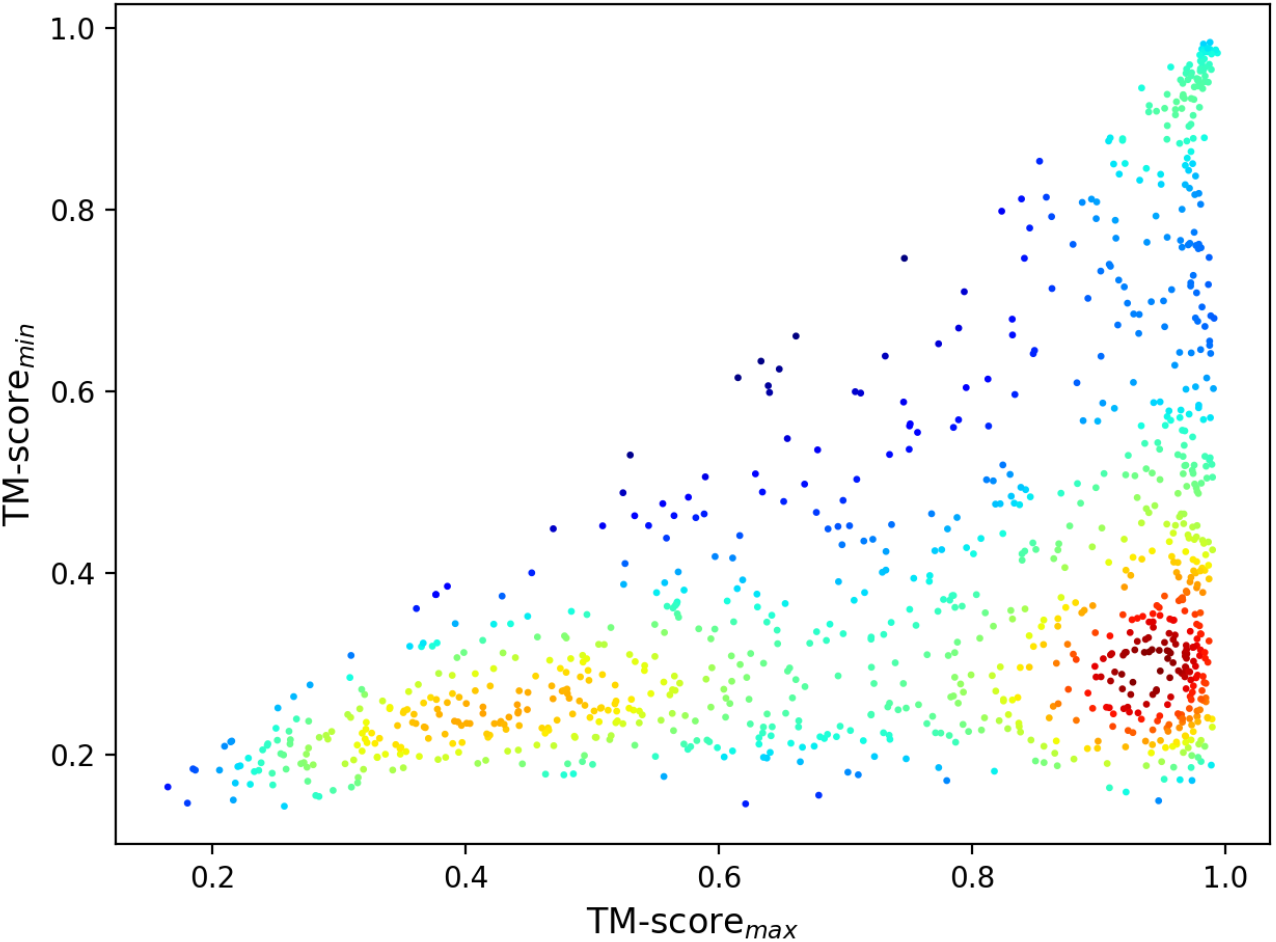
The highest (TM-score_max_) and lowest (TM-score_min_) accuracy values of outputs of ConforFold obtained on dataset B3-2. The plot in is colored by point density.

**Supplementary Figure 11.**
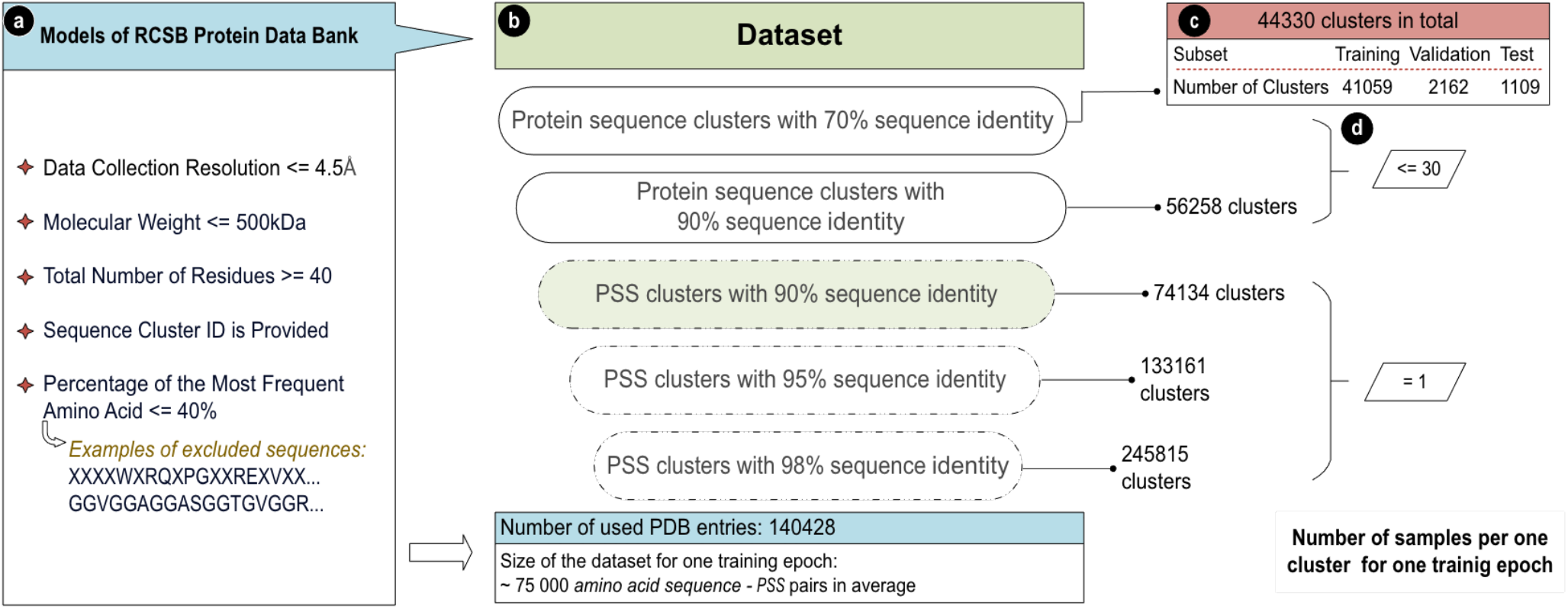
The dataset used in the study for training of Confor-PSSP. **a** Criteria for selecting structures from RCSB PDB. **b** Organization of the dataset. **c** Number of clusters in training, validation and test subsets. **d** Number of samples randomly selected from higher-level clusters during training.

### Supplementary Tables

**Supplementary Table 1.** Performance of ConforPSSP method for 3-state and 8-state PSSP on the test sets at Y=15.

| Test Set | Q3, % | SOV3, % | Q8, % | SOV8, % |
| --- | --- | --- | --- | --- |
| TS | 92.3 | 90.1 | 87.8 | 87.2 |
| CASP10 | 89.8 | 84.4 | 87.1 | 83.6 |
| CASP11 | 88.8 | 82.4 | 86.0 | 81.2 |
| CASP12 | 79.4 | 77.6 | 76.2 | 75.4 |
| CASP13 | 86.0 | 77.2 | 84.0 | 82.1 |

**Supplementary Table 2.**
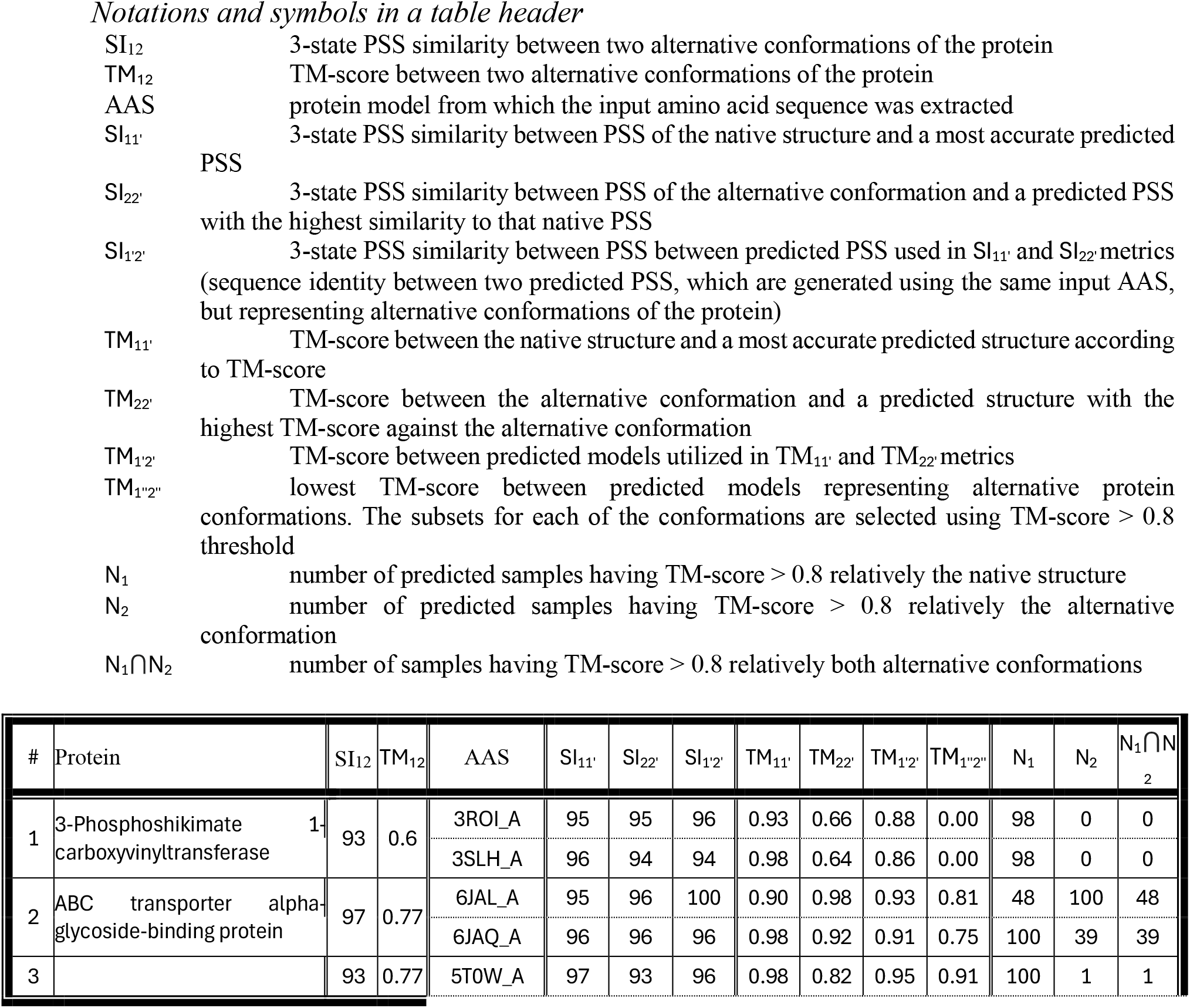
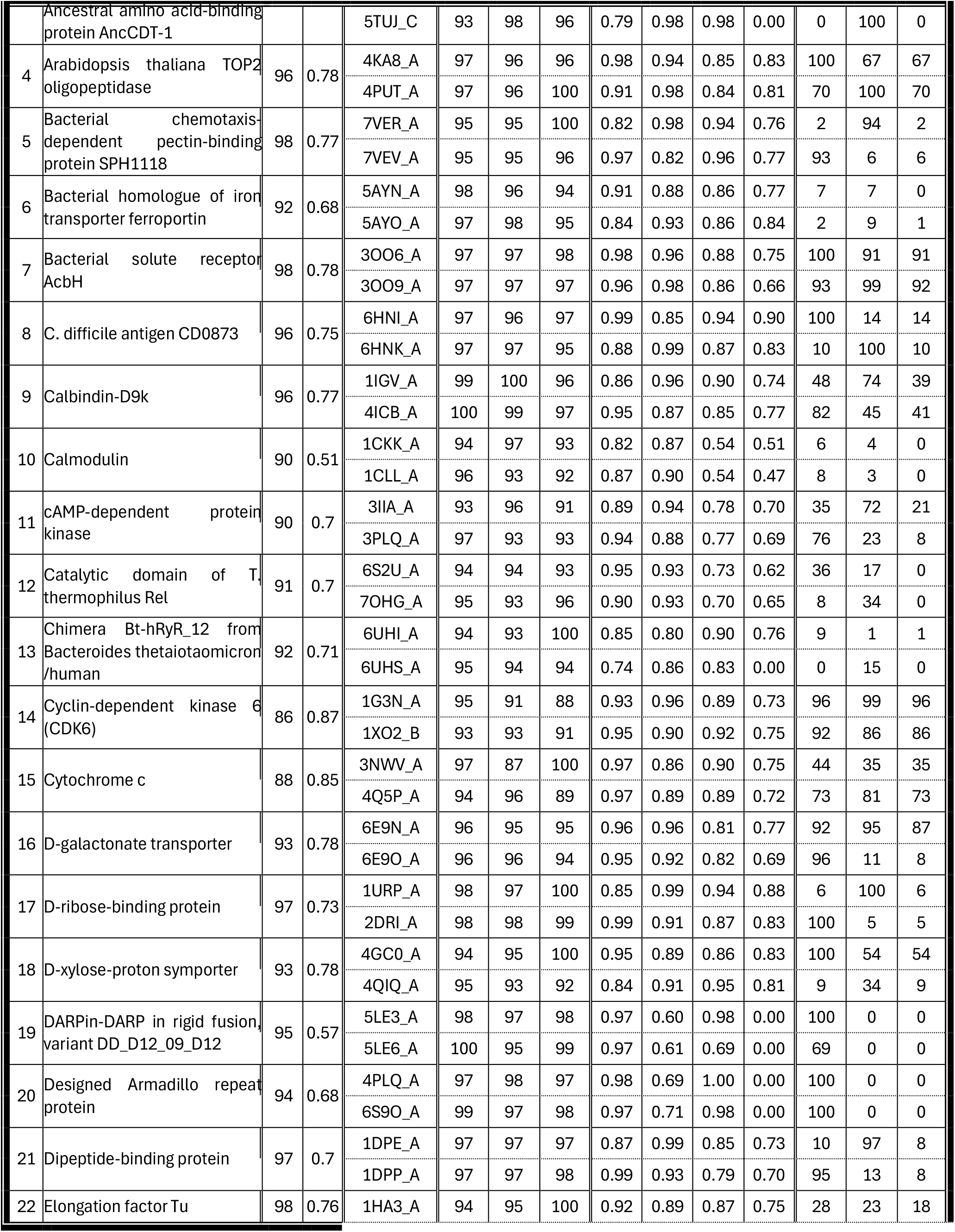
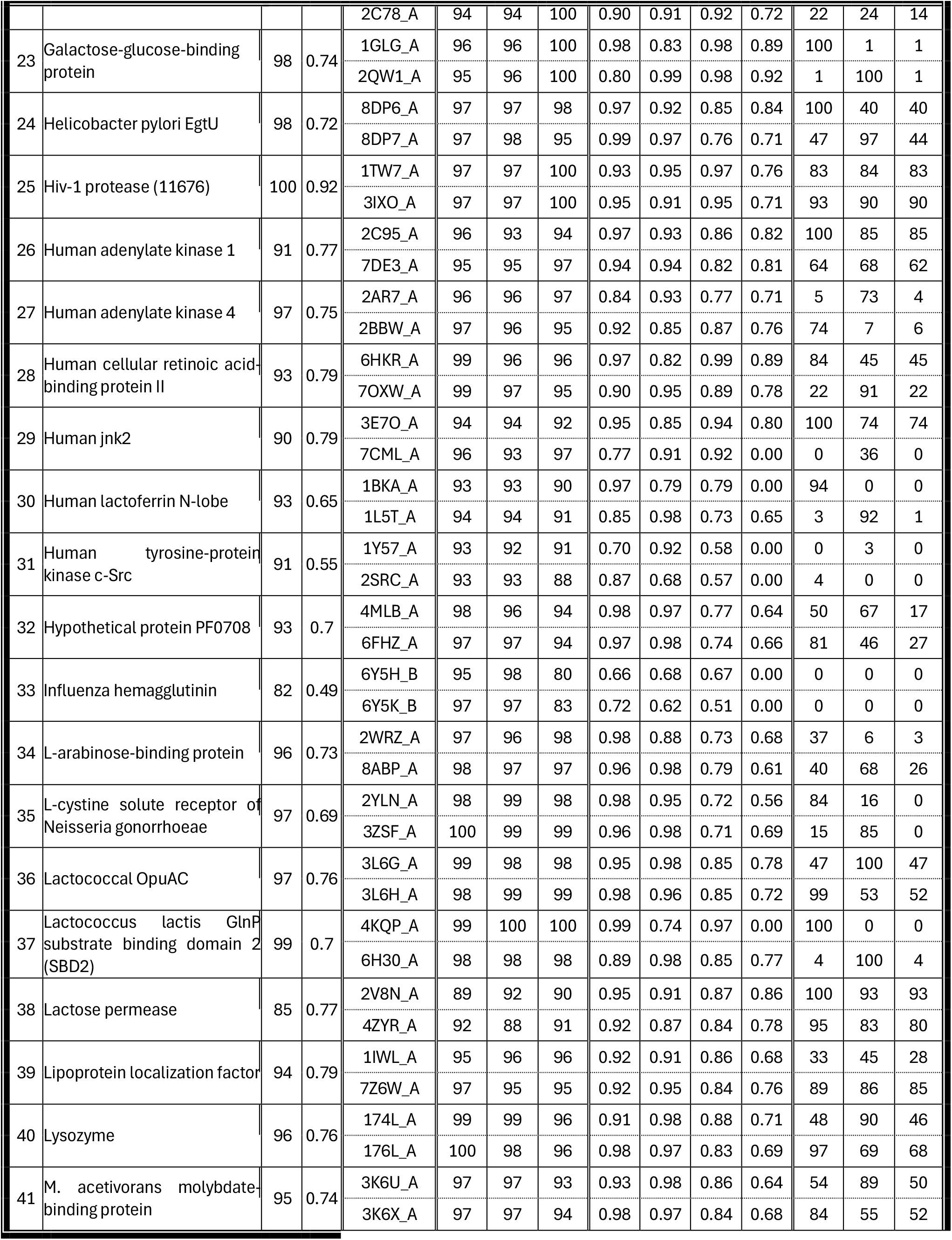
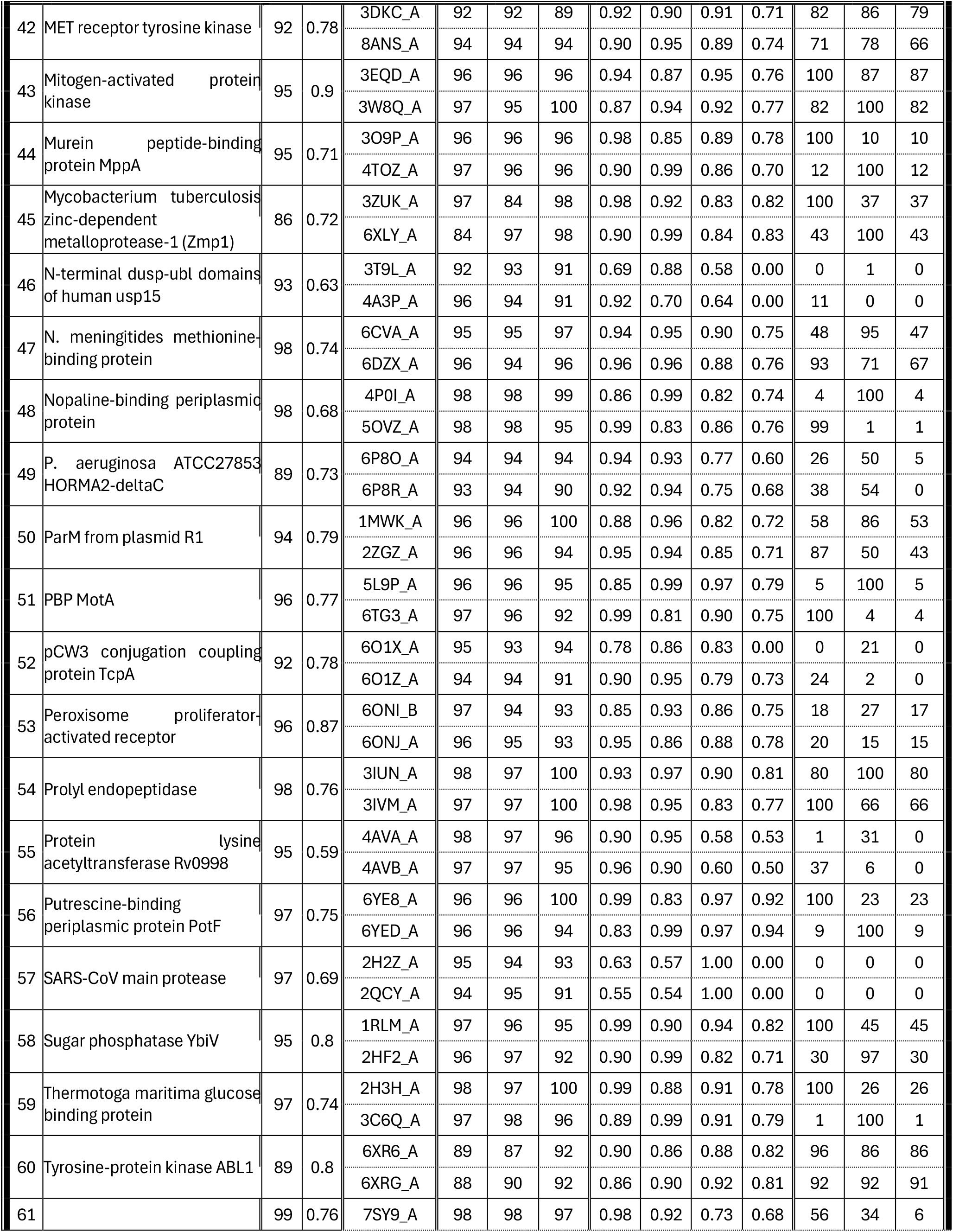
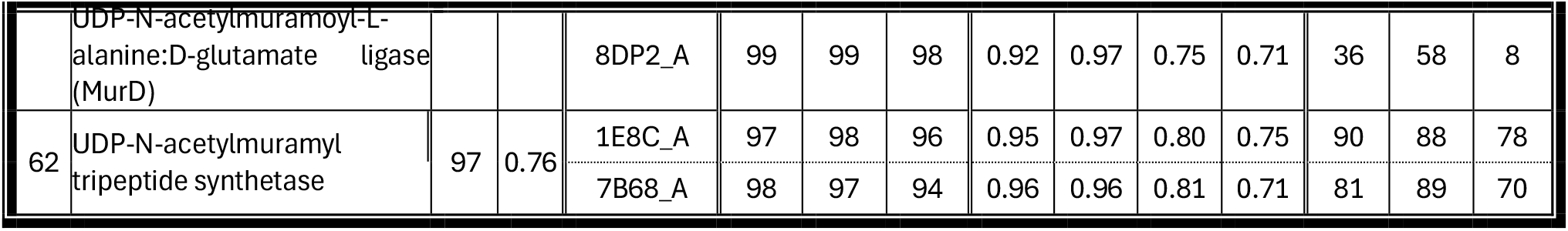
Performance of ConforPSSP and ConforFold methods on prediction of two alternative conformations of proteins from dataset B1.

**Supplementary Table 3.** List of amino acid substitutions.

| *One-letter code of initial amino acid* | *One-letter code of substitute amino acid* | *One-letter code of initial amino acid* | *One-letter code of substitute amino acid* |
| --- | --- | --- | --- |
| B | D | Q | E |
| M | L | U | C |
| N | D | Z | E |

**Supplementary Table 4.** Encoding of the “NWKGIAAMAKKLLXI” sequence.

| Starting residue | Split sequence | Encoded sequence |
| --- | --- | --- |
| 1 | XXXD-WKGI-AALA-KKLL | 1, 97728, 89347, 166, 48892, 2 |
| 2 | XXDW-KGIA-ALAK-KLLX | 1, 97488, 47879, 2924, 49223, 2 |
| 3 | XDWK-GIAA-LAKK-LLXX | 1, 93236, 31083, 52084, 55146, 2 |
| 4 | DWKG-IAAL-AKKL-LXXX | 1, 16420, 40376, 2746, 57397, 2 |

**Supplementary Table 5.** CASP archive files from CASP repository.

| Dataset | Archive |
| --- | --- |
| CASP2010 | casp10.domains_official.noT0695T0739.tgz |
| CASP2011 | casp11.domains_official.release11242014.tgz |
| CASP2012 | casp12.targets_TR.tgz |
| CASP2013 | casp13.targets.ASNX.4public.tar.gz |

**Supplementary Table 6.**
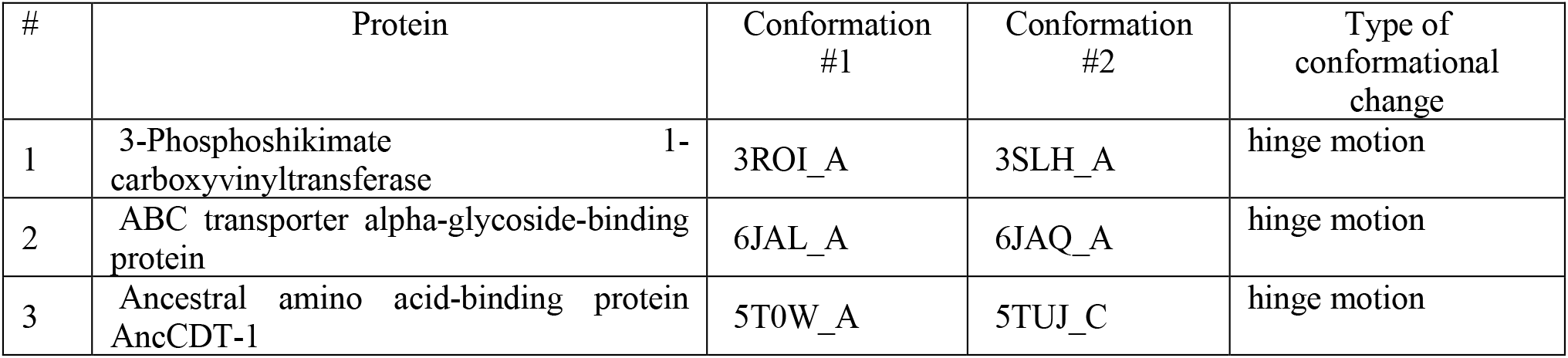
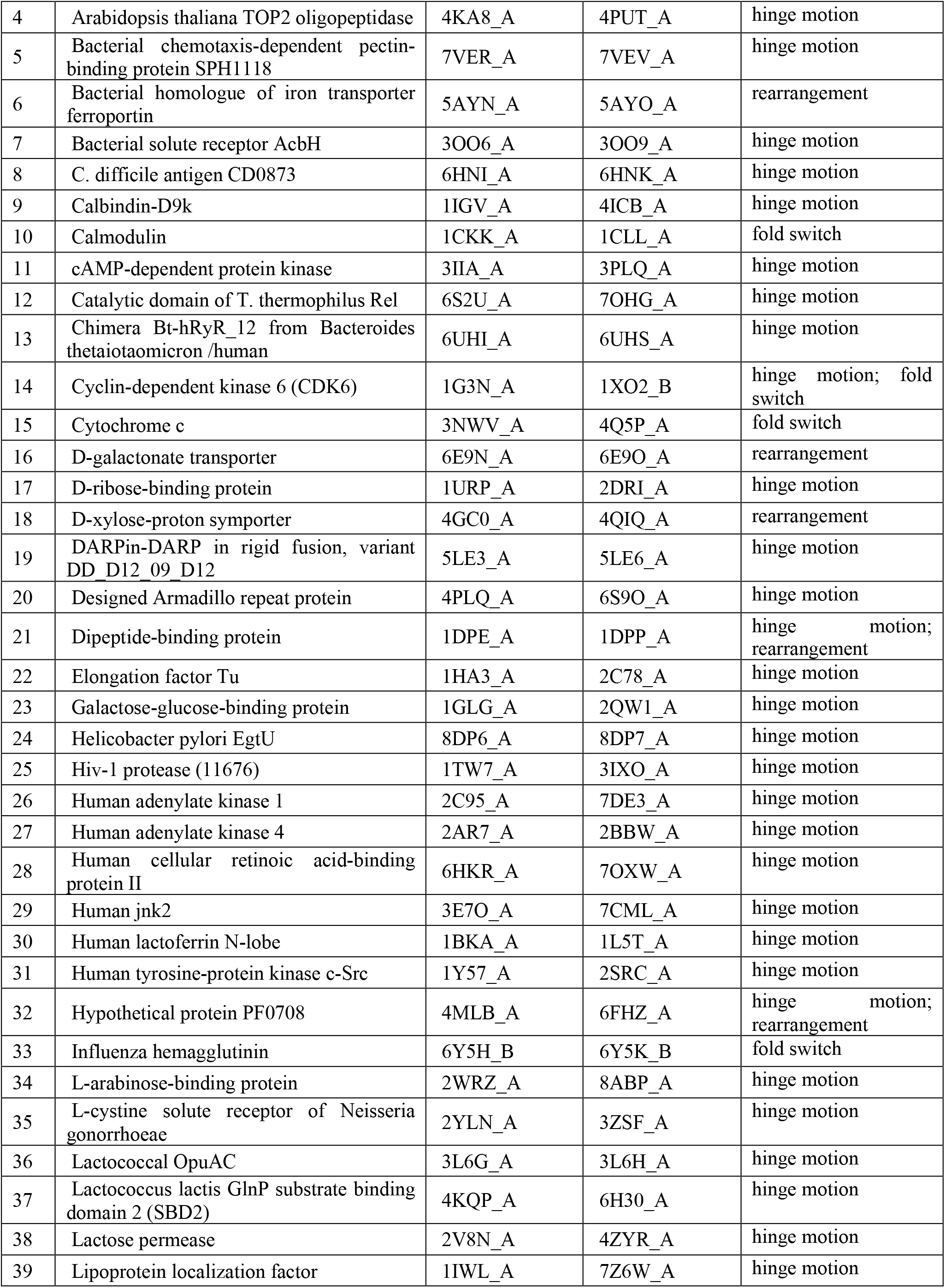
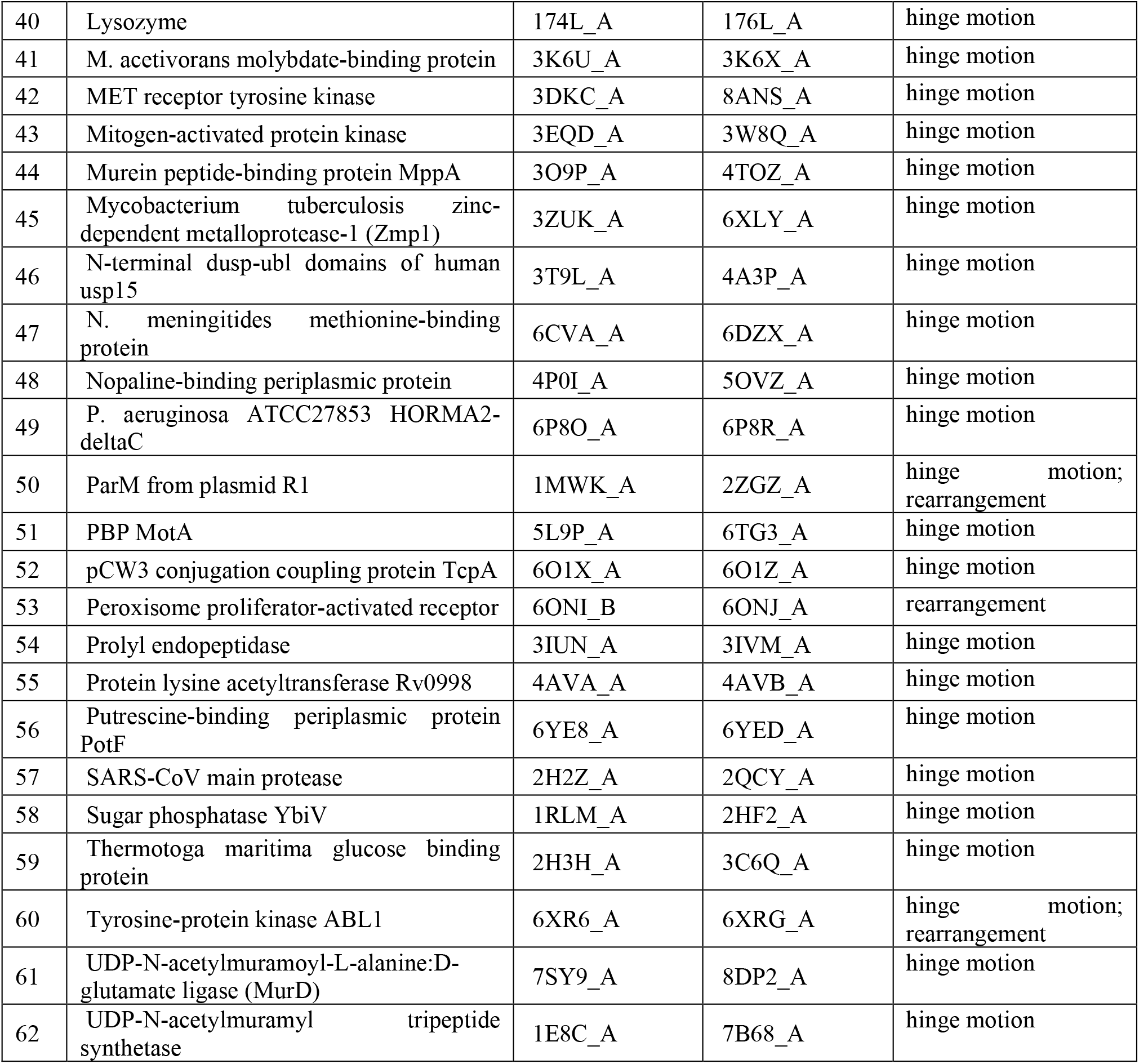
Dataset B1.

*Additional models used in Figure 3*

Calmodulin: 1CKK, 1CLL, 1DMO, 1IQ5, 1NWD, 1PRW, 1X02, 2L53, 2M0K

Cytochrome c: 1HRC, 3NWV, 4Q5P

**Supplementary Table 7.**
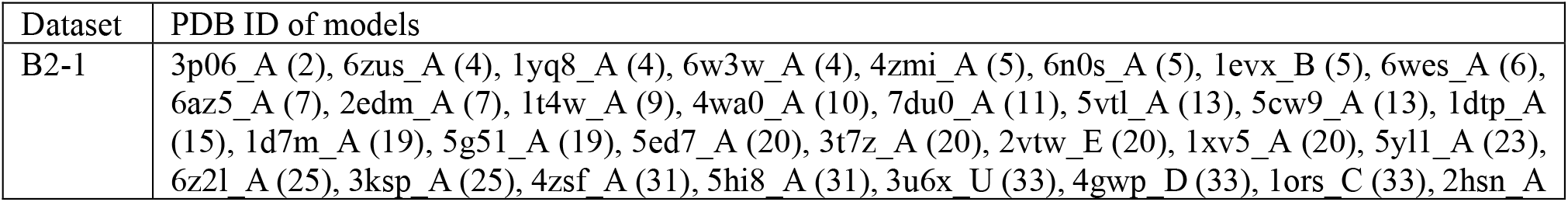
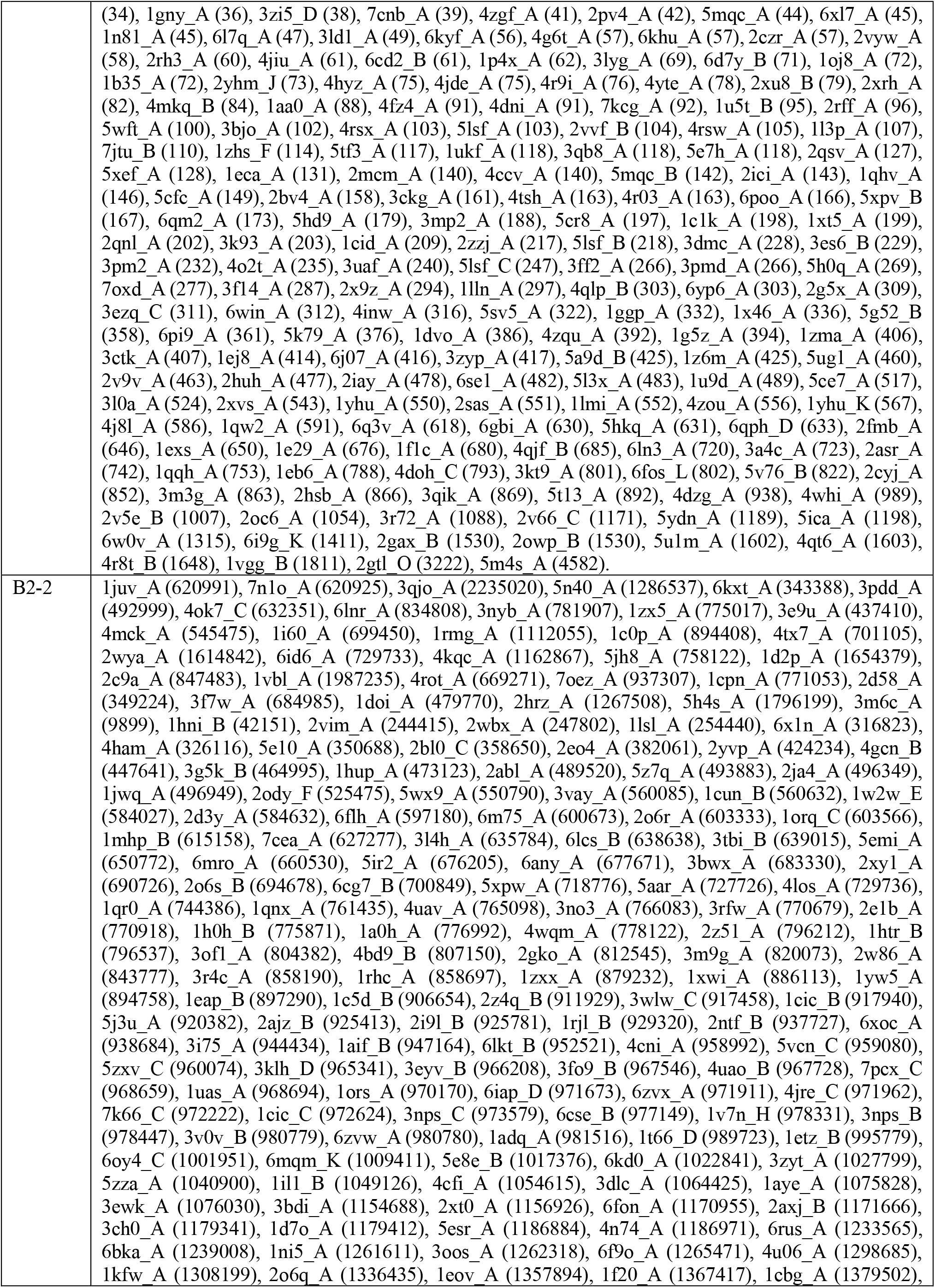
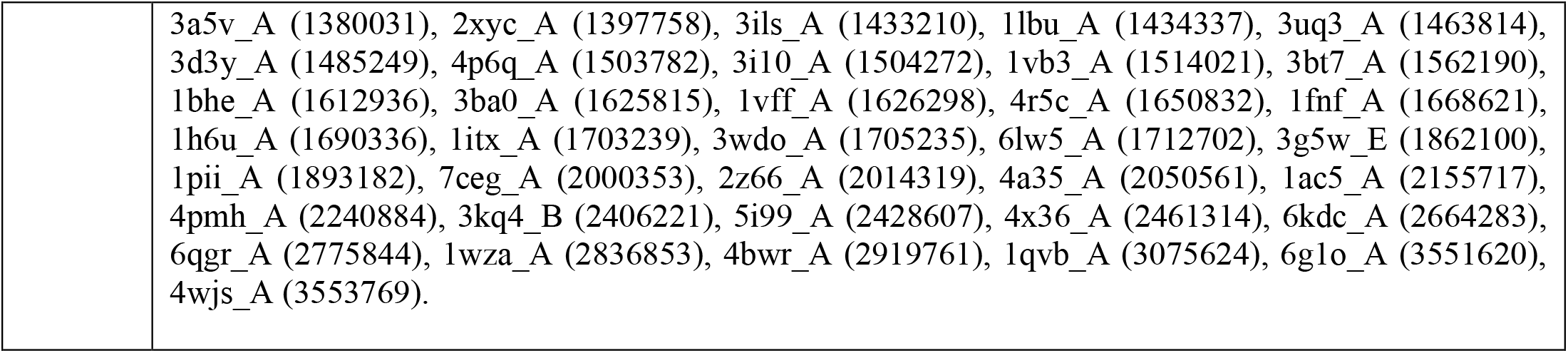
Datasets B2-1 and B2-2. Numbers in parentheses in the second column are number of sequences in MSA and number of gaps in MSA in case of datasets B2-1 and B2-2, respectively.

